# The gut microbiota is associated with clearance of *Clostridium difficile* infection independent of adaptive immunity

**DOI:** 10.1101/512616

**Authors:** Jhansi L. Leslie, Kimberly C. Vendrov, Matthew L. Jenior, Vincent B. Young

## Abstract

*Clostridium (Clostridioides) difficile,* a Gram-positive, anaerobic bacterium is the leading single cause of nosocomial infections in the United States. A major risk factor for *C. difficile* infection (CDI) is prior exposure to antibiotics as they increase susceptibility to CDI by altering the membership of the microbial community enabling colonization. The importance of the gut microbiota in providing protection from CDI is underscored by the reported 80-90% success rate of fecal microbial transplants in treating recurrent infection. Adaptive immunity, specifically humoral immunity, is also sufficient to protect from both acute and recurrent CDI. However, the role of the adaptive immune system in mediating clearance of *C. difficile* has yet to be resolved. Using murine models of CDI, we found that adaptive immunity is dispensable for clearance of *C. difficile*. However, Random Forest analysis using only 2 members of the resident bacterial community correctly identified animals that would go on to clear the infection with 66.7% accuracy. These findings indicate that the indigenous gut microbiota independent of adaptive immunity facilitates clearance of *C. difficile* from the murine gastrointestinal tract.

**Importance:** *C. difficile* infection is a major cause of morbidity and mortality in hospitalized patients in the United States. Currently the role of the adaptive immune response in modulating levels of *C. difficile* colonization is unresolved. This work suggests that the indigenous gut microbiota is a main factor that promotes clearance of *C. difficile* from the GI tract. Our results show that clearance of *C. difficile* can occur without contributions from the adaptive immune response. This study also has implications for the design of preclinical studies testing the efficacy of vaccines on clearance of bacterial pathogens as inherent differences in the baseline community structure of animals may bias findings.

## Introduction

Human disease due to anaerobic bacterium *Clostridium (Clostridioides) difficile* is a significant cause of morbidity and mortality in the US with an estimated 500,000 cases in the U.S. yearly (1). A major risk factor for *C. difficile* infection (CDI) is prior exposure to antibiotics (2). Antibiotics increase susceptibility to CDI by altering the membership of the microbial community and thus the metabolome of the gut, enabling colonization (3). Colonization with *C. difficile* can manifest in a range of clinical syndromes ranging from asymptomatic colonization to inflammatory colitis characterized by diarrhea with abdominal pain, and in severe cases, death. In addition to primary infection, one in five patients treated for CDI experiences recurrent disease (1).

Disease is primarily mediated by the production of two toxins, TcdA and TcdB, which are the major virulence factors for *C. difficile* (4). TcdA and TcdB are large multi-domain proteins, which inactivate cellular rho-family GTPases via the addition of a glucose molecule (5). Inactivation of these key regulatory proteins in epithelial cells results in disruption of tight junctions, increased paracellular flow, and eventually leads to cell death (6, 7).

The importance of the gut microbiota in providing protection from CDI is underscored by the reported 80-90% success rate of fecal microbial transplants in preventing recurrent infection (8-10). Other than microbiome-mediated prevention of colonization, adaptive immunity is also sufficient to provide protection from both acute and recurrent CDI likely via antibody-mediated neutralization of *C. difficile* toxins TcdA and TcdB (11–14). However, the role of the adaptive immune system in modulating *C. difficile* colonization has yet to be resolved.

In this study we sought to determine if adaptive immunity plays a role in clearance of *C. difficile* colonization. We found clearance of *C. difficile* can occur in the absence of adaptive immunity. Furthermore, the indigenous microbial community membership that exists prior to antibiotic administration and infection was predictive of which animal went on to clear the infection.

## Results

### Clearance of C. *difficile* can occur in the absence of adaptive immune responses

We sought to determine the contribution of adaptive immunity in clearance of *C. difficile*. To test this, we compared *C. difficile* infection in wild-type mice (WT) to RAG1^−/−^ mice, which lack both B and T cells. As the two genotypes of mice were derived from separate colonies and others have reported that RAG1^−/−^ mice have a distinct microbial community from WT mice, we co-housed the RAG1^−/−^ mice with WT mice for over three weeks. Co-housing normalized the WT and RAG1^−/−^ mice fecal communities such that they were not significantly different, ANOSIM *p*=0.087 (figure 1A). Both groups of mice were pretreated with antibiotics, separated into cages based on genotype, then challenged with *C. difficile* strain 630. Although all mice were initially colonized, within three weeks of challenge, animals in two cages cleared *C. difficile* while the remaining animals were persistently colonized (figure 1B). Notably, the mice that cleared the infection were WT and RAG1^−/−^. Clearance or persistence colonization with *C. difficil*e did not correspond to genotype but rather to co-housing group. Reanalyzing the pre-antibiotic microbial communities by co-housing group rather than genotype, we found that the mice that eventually cleared *C. difficile* had significantly distinct community compared to the mice that remained colonized ASOSIM *p*= 0.047 (figure 1C). These results demonstrate that clearance of *C. difficile* can occur independently of adaptive immunity.

### Reconstitution of IgG anti-toxin antibody is not sufficient to clear *C. difficile*

To mitigate any effect inherent baseline differences in the microbiota of WT and RAG1^−/−^ mice, we tested if adaptive immunity is sufficient to clear *C. difficile* by reconstituting RAG1^−/−^ mice with splenocytes from WT mice. Reports of immunization with various *C. difficile* antigen suggests that antibodies to these antigens may decrease colonization so we additionally tested if transfer of cells from mice immunized via natural infection with *C. difficile* might facilitate clearance (15, 16). Splenocytes were collected from WT mice that were either naïve or colonized with *C. difficile* strain 630 for three weeks (figure S1A). Development of humoral immune responses to *C. difficile* in the donor mice was confirmed by the detection of high titers of anti-TcdA IgG in the serum while uninfected mice had undetectable levels of anti-TcdA serum IgG (*p*<0.01) (figure S1B).

Recipient RAG1^−/−^ mice were infected with *C. difficile* strain 630 prior to the adoptive transfer. Donor splenocytes were administered to the recipient RAG1^−/−^ mice two days after *C. difficile* challenge, when *C. difficile* colonization had already reached high levels. Recipient mice were randomly assigned to one of three groups and either received splenocytes from naïve WT donors, infected WT donors, or vehicle.

To confirm engraftment of the WT cells, we measured total serum IgG in the recipient mice three-weeks post transfer. The mice that received splenocytes had significantly higher levels of total serum IgG post-transfer compared to the mice that received vehicle (*p*<0.05) (figure 2A). Of the mice that received splenocytes, two did not develop any detectable serum IgG. There was no difference in the levels of total serum IgG between the mice that received splenocytes from infected donors versus uninfected donors (*p*>0.05). Furthermore, we determined that we successfully transferred anti-*C. difficile* immunity as we detected anti-TcdA IgG only in the serum from the mice that received splenocytes from the infected donors (*p*<0.01) (figure 2B).

Following adoptive transfer, levels of *C. difficile* in the feces were monitored for three weeks. We observed clearance of *C. difficile* from one cage of mice in the group that received splenocytes from infected donors. However, clearance of *C. difficile* did not occur in any of the other animals within that treatment group (figure 2C). Three-weeks post transfer there was no significant difference in levels of colonization in any of the treatment groups (figure 2D). Notably, in the cage that cleared, one mouse had undetectable levels of serum IgG while the other three mice in the cage had detectable levels (figure 2A, filled pink circles). Together these results suggest that reconstitution of adaptive immunity is not sufficient for clearance of *C. difficile*.

The range in the levels of colonization we observed within each treatment group suggested adaptive immunity is not sufficient to explain the differences in clearance of *C. difficile*. Visualization of the Bray-Curtis dissimilarity between the day one post infection communities (before the adoptive transfer) using multidimensional scaling revealed that the mice that went on to clear *C. difficile* had a distinct community compared to the mice that would remain colonized, ANOSIM *p*=0.02 (figure 3). This result suggests the structure of gut microbiota rather than adoptive transfer of splenocytes is associated with clearance of *C. difficile*. \

### Specific members of the microbiota are altered in mice with reconstituted adaptive immunity

The microbiota and the immune system have been previously shown to modulate one another through numerous complex interactions (18, 19). In the cefoperazone mouse model of infection, the diversity of microbiota begins to recover by two weeks following cessation of the antibiotic (20). Therefore, we asked if reconstitution of adaptive immunity altered the recovery of the community following antibiotics and infection with *C. difficile*. We examined the gut microbial community structure of the mice over the course of the experiment using 16S rRNA gene amplicon sequencing. Our first approach sought to determine if we could detect changes in the overall microbial community composition of the mice. We calculated the Bray-Curtis dissimilarity between each mouse’s day twenty-one sample (nineteen days after the adoptive transfer) and their pre-antibiotic sample. We hypothesized that reconstitution of adaptive immunity might prevent the microbiota from returning to the same structure as was observed before adoptive transfer. Thus, we thought that perhaps the mice that received splenocytes might have higher Bray-Curtis dissimilarity values compared to the group that received only vehicle. Since we were unable to confirm that we successfully restored adaptive immune function in two of mice that received splenocytes (figure 2A), we excluded them from the rest of analysis as our questions hinged on immune status-gut microbiota interactions. Additionally, we lost the ability to calculate this metric for a couple of mice due to the lack of pre-antibiotic samples. Comparing the Bray-Curtis dissimilarity results between the three treatment groups revealed no significant differences between any of the groups (figure 4A). We also wondered if addition of adaptive immunity might alter alpha-diversity so we calculated the inverse Simpson index each fecal community at day nineteen post transfer (day twenty-one post infection). We did not observe any significant differences between the treatment groups by this metric either (figure 4B). This suggested that by broad evaluations of community structure, the perturbation of antibiotics and infection with *C. difficile* potentially has a much greater effect on the microbial community than any effects due to immune reconstitution.

While we saw no significant differences in the recovery of the community structure or alpha diversity at day twenty-one post infection, we wondered if perhaps the abundance of only a few operational taxonomic units (OTUs) were altered by reconstitution of the adaptive immune system. For this analysis, we grouped all of the mice that received splenocytes and developed detectable levels of serum IgG at day twenty-six post infection together and called them IgG positive. The mice that only received vehicle and thus had undetectable levels of serum IgG were designated the IgG negative group. Using OTU abundance from day twenty-one post infection samples, linear discriminant analysis effect size (LefSe) identified twenty-seven OTUs with linear discriminant analysis (LDA) values greater than two. The ten OTUs with the highest LDA values were primarily enriched in the IgG negative mice (figure 4C). OTU 3, which is classified as *Akkermansia*, had the highest LDA value. This OTU was found at a significantly lower abundance in the IgG positive mice compared to the IgG negative mice. A decrease in *Akkermansia* following reconstitution of adaptive immunity via transfer of bone marrow of from wild-type mice into RAG1^−/−^ mice has been reported by another group (21). While the decrease in OTU 3 in the IgG positive mice was observed across the cages, many of the other OTUs that discriminated between the IgG positive and negative mice were only detected in one of the IgG negative cages.

### Random Forest feature selection identifies OTUs in the pre-antibiotic community that differentiates mice that will remain persistently colonized vs. clear

Following our previous analyses, we made the consistent observation that structure of the gut microbiome was associated with clearance of *C. difficile*, even prior to antibiotic treatment (figure 1C). We questioned if specific OTUs present in the mice before any intervention may have differentiated mice that would go on to clear the infection. For this analysis, we pooled data from three independent experiments (the two described earlier and a third experiment including only WT mice) where cages of mice had spontaneously cleared *C. difficile* (figure S2). We utilized Random Forest for feature selection to identify OTUs that could classify mice as “cleared” or “colonized” based on their pre-intervention microbiota. Using the entire pre-treatment community, we could classify the mice as “cleared” or “colonized” with 76.9% accuracy. However, this model was better at classifying mice that would remain colonized and was poor at classifying mice that would go onto clear *C. difficile*, with an accuracy of only 25%. Nine out of the top ten OTUs that most contributed to classification were from the Firmicutes phylum (figure S3A and B). Two OTUs in particular (OTUs 52 and 93) ranked highest in their ability to discriminate between the groups and were significantly increased in abundance in mice that would go on to clear. Therefore, we tested if those two OTUs alone were sufficient to classify the mice. Generating a new Random Forest model using only those two OTUs, we found that the overall model improved to 82.9% accuracy in classification. Furthermore, these two OTUs could correctly classify mice that would go on to clear *C. difficile* with 66.6% accuracy (Figure 5).

## Discussion

In this study we asked if adaptive immunity was required for clearance of the gastrointestinal pathogen *C. difficile*. Results from multiple experimental models lead us to conclude that clearance of *C. difficile* in mice can occur without contributions from adaptive immune responses. This finding is in contrast to the paradigm observed in other gastrointestinal infections. For example, infection with the attaching-effacing pathogen *Citrobacter rodentium*, provides a framework by which the adaptive immunity facilitates clearance (22, 23). In addition to the potential direct effects, adaptive immunity may have on the bacterium itself, it is known that there is a complex interaction loop between the microbiota and host immune response. Both the innate and adaptive arms of the immune system regulate membership of the gut microbial community while the gut microbiota in turn modulates the immune system via the production of metabolites and/or MAMPs (24).

Our results show that reconstitution of adaptive immunity is associated with altered abundance of some bacteria in the gut, however it does not impact levels of *C. difficile* colonization. We found that in the reconstituted RAG1^−/−^ mice that developed serum IgG, there was a decreased abundance of *Akkermansia* (OTU 3). Another group has previously observed this result; however, we were surprised to see the same trend in our model as our mice were also subjected to antibiotic therapy and infection with *C. difficile.* In the two mice that received splenocytes but did not have detectable serum IgG, the abundance of the *Akkermansia* OTU was very low (figure S4A). There are numerous reasons why this could be the case, the first being that a lack of serum IgG does not preclude successful transfer of T cells which may be responsible for modulating levels of *Akkermansia* in wild-type mice. Additionally, fecal IgG or IgA from the mice that had successful transfers may have been transmitted via coprophagy in sufficient quantities to modulate the levels of *Akkermansia* in the IgG negative mice that were sharing their cage. Since the relative abundance of OTU3 was not significantly different between the groups in the pre-treatment samples we can conclude that the differences we observed were a result of the experimental conditions not merely baseline differences in their microbiota (figure S4B). *Akkermansia* has been implicated in the modulation of health processes such as regulation of host metabolism so further studies are necessary to fully elucidate the factors that regulate its abundance in the gut (25, 26).

Based on our repeated observations that altered communities early in the experimental timeline was associated with clearance of *C. difficile* we used Random Forest to eventually identify just two OTUs that could classify mice that would go on to clear C. *difficile* with 66.6% accuracy. Previous work using a similar approach identified OTUs present on the day of challenge that were predictive of levels of colonization day one post-infection, however we are the first group to assess if the composition of the murine gut microbiota before any treatment might affect the outcome of C. *difficile* infection (27). Both of the OTUs we identified belong to the family *Lachnospiraceae* and were enriched in mice that would go on to clear *C. difficile* infection. Our group has previously observed that high levels of *Lachnospiraceae* is associated with protection from severe disease in a murine model of CDI (28). One possibility is that these bacteria are just inherently resistant to cefoperazone, however in vitro antibiotic susceptibility testing of *Lachnospiraceae* isolates from our mouse colony suggest that this is not the case (data not shown). Furthermore, we have also reported that mono-association of germ-free mice with a single *Lachnospiraceae* isolate partially restored colonization resistance (29). It is tempting to speculate multiple *Lachnospiraceae* isolates might be able to fully restore colonization resistance. However, it remains to be seen if the same mechanisms, which prevent initial colonization of *C. difficile*, play a role in clearance of *C. difficile.*

Our results suggest that community resilience is intrinsic to the community membership at baseline, prior to any antibiotic treatment. Additionally, these data suggest the possibility of predicting individuals that will be at risk for persistent colonization before antibiotic therapy. However, a crucial first step is to determine if predictive OTUs are different across perturbations such as various classes of antibiotic therapy. Finally, our findings have implications for the design of future preclinical studies testing the efficacy of vaccines or other manipulations of adaptive immunity on levels of colonization as “cage effects” or inherent differences in the baseline community structure of animals within cages may bias findings. Experimental approaches that can be implemented to account for the role of the microbiota include co-housing, using multiple cages for each experimental condition, and the use of littermate controls (30).

## Material and Methods

### Animal Husbandry

Both male and female C57BL/6 specific-pathogen-free (SPF) mice age five to twelve weeks were used in these studies. The wild-type (WT) mice were from a breeding colony at the University of Michigan, originally derived from Jackson Laboratories over a decade ago. The RAG1^−/−^ (B6.129S7-*Rag1*^*tm1Mom*^/J) mice were from a breeding colony started with mice from Jackson Laboratories in 2013. Animals were housed in filter top cages with corncob bedding and nestlet enrichment. Water bottles were autoclaved empty and filled in a biological safety cabinet with either sterile water or antibiotic dissolved in sterile water. Mice were fed a standard irradiated chow (LabDiet 5LOD) and had access to food and water *ad libitum*. Cage changes were carried out in a biological safety cabinet. The frequency of cage changes varied depending on the experiment. To prevent cross-contamination between cages, hydrogen peroxide-based disinfectants in addition to frequent glove changes were utilized during all manipulation of the animals. The mice were maintained under 12-hours of light/dark cycles in facilities maintained at a temperature of 72°C +/− 4 degrees. Animal sample size was not determined by a statistical method. Multiple cages of animals for each treatment were used to control for possible differences in the microbiota between cages. Mice were evaluated daily for signs of disease. Euthanasia was carried out via CO_2_ asphyxiation when mice were determined to be moribund or at the conclusion of the experiment. Animal studies were conducted under the approval of The University of Michigan Committee on the Care and Use of Animals; husbandry was performed in an AAALAC-accredited facility.

### Spore Preparation

Spore stocks of *C. difficile* strain 630 (ATCC BAA-1382) were prepared as previously described with the following modifications; strains were grown overnight in 5mL of Difco Columbia broth (BD Biosciences 294420), which was added to 40mL of Clospore media (3, 31).

### Infections

In experiments comparing colonization in WT and RAG1^−/−^ mice, age and sex matched mice were co-housed for thirty-three days starting at three weeks of age and continuing through cefoperazone administration. Upon infection, animals were separated into single genotype housing. Mice were made susceptible to infection by providing *ad libitum* drinking water with the addition of 0.5mg/mL cefoperazone (MP Pharmaceuticals 0219969501) in Gibco distilled water (15230147). The antibiotic water was changed every two days and was provided for ten days. Following two days of supplying drinking water without antibiotic, mice were challenged with either spores or water (mock). *C. difficile* spores suspended in 50μL of Gibco distilled water were administered via oral gavage. The number of viable spores in each inoculum was innumerate by plating for colony-forming units (CFU) per mL^−1^ on pre-reduced taurocholate cycloserine cefoxtin fructose agar (TCCFA). TCCFA was made as originally described (32) with the following modifications: the agar base consisted of 40g of Proteose Peptone No. 3 (BD Biosciences 211693), 5g of Na_2_HPO_4_ (Sigma-Aldrich S5136), 1g of KH_2_PO_4_ (Fisher P285500), 2g NaCl (J.T. Baker 3624-05), 0.1g MgSO_4_ (Sigma M7506), 6g Fructose (Fisher L95500), and 20g of agar (Life Technologies 30391-023) dissolved in 800mL of Milli-Q water. Following adjustment of volume to 1L, the media was autoclaved and supplemented with D-cycloserine to a final concentration of 250μg/mL of (Sigma-Aldrich C6880), cefoxitin to a final concentration 16μg/mL (Sigma-Aldrich C4786) and taurocholate to a final concentration 0.1% (Sigma T4009). Over the course of the infection, mice were routinely weighed and stool was collected for quantitative culture. Mice were challenged with between 10^2^ and 10^4^ CFU.

### Quantitative Culture

Fresh voided fecal pellets were collected from each mouse into a pre-weighted sterile tube. Following collection, the tubes were reweighed and passed into an anaerobic chamber (Coy Laboratories). In the chamber, each sample was diluted 1 to 10 (w/v) using pre-reduced sterile PBS and serially diluted. 100μL of a given dilution was spread onto pre-reduced TCCFA or when appropriate TCCFA supplemented with final concentration of either 2 or 6μg/mL of erythromycin (Sigma E0774). Strain 630 is erythromycin resistant; use of erythromycin in TCCFA plates reduced background growth from other bacteria in the sample. Plates were incubated anaerobically at 37°C and colonies were enumerated at 18-24 hours. Plates that were used to determine if mice were negative for *C. difficile* were held and rechecked at 48 hours.

### Splenocytes Recovery and Transfer

Spleens from individual animals were aseptically harvested from donor mice. Following harvest, the organ was gently homogenized using sterile glass slides to remove the cells from the capsule. Cells were suspended in filter-sterilized RPMI complete media consisting of RPMI + 1 L-glutamine (Gibco 11875-093) supplemented with 10% FBS (Gibco 16140-071), 1% 100x Penicillin-Streptomycin (Gibco 15070-063), 1% 1M HEPES (Gibco 15630-080), 1% 100x non-essential amino acids (Gibco 11140-050), 1% 100mM Sodium Pyruvate (Gibco 11360-070) and 0.05mL of 1M 2-Mercaptoethanol (Sigma M3148). To remove large debris, the cell suspension was filtered through a 40μm cell strainer. Cells were pelleted by centrifugation at 1,500 rpm for 5 minutes at 4°C. Following the spin, the pellet was suspended in red blood cell lysing buffer (Sigma R7757) and incubated with the solution for no more than 5 minutes. Lysis was stopped with the addition of RMPI complete media and cells were enumerated manually using a haemocytometer. Following enumeration, the cells were pelleted again by centrifugation at 1,500 rpm for 5 minutes at 4°C and re-suspended in Leibovitz’s L-15 (Corning 10-045-CV) media. Recipient mice were injected into the peritoneal cavity with 2 × 10^7^ cells in 0.25mL L-15 media. Mice that received vehicle were injected with 0.25mL of L-15 media only.

### Blood Collection

Blood was collected from either the saphenous vein for pre-treatment time points or via heart puncture at the experimental endpoint. Collections from the saphenous vein utilized capillary tubes (Sarstedt microvette CB300 Z) while blood collected via heart puncture utilized a polymer gel-based separator tube (BD Microtainer SST). Following collection, tubes were spun according to manufacturer’s instructions, serum was aliquoted and stored at −80°C until use.

### Total IgG ELISA

Total serum IgG levels were measured using the IgG (Total) Mouse Uncoated ELISA Kit (ThermoFisher Scientific 88-50400). Each sample was diluted 500-fold in assay buffer and run in duplicate with Southern Biotech TMB Stop Solution (0412-01) used as the stop solution. Optical density values were measured at 450nm and 570nm on a VersaMax plate reader (Molecular Devices, Sunnyvale, CA) and corrected by subtracting the 570nm measurement from the 450nm measurement. A 4-Parameter Standard Curve was used to calculate sample concentration values.

### Anti-*C. difficile* TcdA IgG ELISA

Titers of serum IgG specific to *C. difficile* TcdA (toxin A) was measured by ELISA as previously described, with the following modifications (33). Serum from RAG1^−/−^ mice that received an adoptive transfer was diluted 1:50 in blocking buffer with subsequent serial dilutions to a final dilution of 1:12,150. Serum from the wild-type mice was diluted 1:1,200 in in blocking buffer with subsequent serial dilutions to a final dilution of 1:874,800. Each sample was run in duplicate. Each plate had the following negative controls: all reagents except serum, all reagents except toxin and pre-immune serum if applicable. Additionally, each plate had a positive control consisting of toxin coated wells reacted with mouse TcdA monoclonal antibody TGC2 diluted 1:5,000 in blocking buffer (antibodies-online.com ABIN335169). The optical density at 410nm and 650nm was recorded on a VersaMax plate reader (Molecular Devices, Sunnyvale CA). The absorbance for each sample was corrected by subtracting the OD_650_ reading from the OD_410_ reading. The anti-TcdA IgG titer for each sample was defined as the last dilution with a corrected OD_410_ greater than average corrected OD_410_ of the negative control wells plus three times the standard deviation of those wells.

### DNA Extraction

Genomic DNA was extracted from approximately 200-300μl of fecal sample using the MoBio PowerSoil HTP 96 DNA isolation kit (formerly MoBio, now Qiagen) on the Eppendorf EpMotion 5075 automated pipetting system according to manufacturer’s instructions.

### Sequencing

The University of Michigan Microbial Systems Laboratory constructed amplicon libraries from extracted DNA as described previously (34). Briefly, the V4 region of the 16S rRNA gene was amplified using barcoded dual index primers as described by Kozich et al. (35). The PCR reaction included the following: 5μl of 4μM stock combined primer set, 0.15μl of Accuprime high-fidelity Taq with 2μl of 10× Accuprime PCR II buffer (Life Technologies, 12346094), 11.85μl of PCR-grade water, and 1μl of template. The PCR cycling conditions were as follows: 95°C for 2 minutes, 30 cycles of 95°C for 20 seconds, 55°C for 15 seconds, and 72°C for 5 minutes, and 10 minutes at 72°C. Following construction, libraries were normalized and pooled using the SequelPrep normalization kit (Life Technologies, A10510-01). The concentration of the pooled libraries was determined using the Kapa Biosystems library quantification kit (KapaBiosystems, KK4854) while amplicon size was determined using the Agilent Bioanalyzer high-sensitivity DNA analysis kit (5067-4626). Amplicon libraries were sequenced on the Illumina MiSeq platform using the MiSeq Reagent 222 kit V2 (MS-102-2003) (500 total cycles) with modifications for the primer set. Illumina’s protocol for library preparation was used for 2nM libraries, with a final loading concentration of 4pM spiked with 10% genomic PhiX DNA for diversity. The raw paired-end reads of the sequences for all samples used in this study can be accessed in the Sequence Read Archive under PRJNA388335.

### Sequence Curation and Analysis

Raw sequences were curated using the mothur v.1.39.0 software package (36) following the Illumina MiSeq standard operating procedure. Briefly, paired end reads were assembled into contigs and aligned to the V4 region using the SLIVA 16S rRNA sequence database (release v128) (37), any sequences that failed to align were removed; sequences that were flagged as possible chimeras by UCHIME were also removed (38). Sequences were classified with a naïve Bayesian classifier (39) using the Ribosomal Database Project (RDP) and clustered in to Operational Taxonomic Units (OTUs) using a 97% similarity cutoff with the Opticlust clustering algorithm (40).

The number of sequences in each sample was then rarefied to 10,000 sequences to minimize bias due to uneven sampling. For feature selection, the shared file was filtered to remove any OTU that was in less than six samples across the entire data set. The mothur implementation of LefSe (linear discriminant analysis effect size) was used to determine OTUs that differentiated IgG positive verses RAG1^−/−^ mice given vehicle nineteen days post adoptive transfer (41). Following curation in mothur, further data analysis and figure generation was carried out in R (v 3.3.3) using standard and loadable packages (42). The data and code for all analysis associated with this study are available at https://github.com/jlleslie/AdaptiveImmunity_and_Clearance.

Most of the analysis relied on the R package vegan (43). This includes, determining the axes for the multidimensional scaling (MDS) plots using Bray-Curtis dissimilarity calculated from sequence abundance. Additionally, vegan was used to determine significance between groups using ANOSIM, calculation of Inverse Simpson index, and Bray-Curtis dissimilarity between samples. Final figures were modified and arranged in Adobe Illustrator CC. For the purpose of distinguishing between values that were detected at the limit of detection versus those that were undetected, all results that were not detected by a given assay were plotted at an arbitrary point below the LOD. However, for statistical analysis, the value of 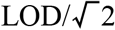 was substituted for undetected values. Wilcoxon ranked sum test was used to determine significant differences and when appropriate, reported *p*-values were corrected for multiple comparisons using the Benjamini–Hochberg correction.

### Random Forest Analysis

Random Forest analysis was performed using R (v.3.2.3) using the randomForest package (44, 45). Model parameters ntree and mtry were tuned based on the input datasets in order to achieve optimal classification without over-fitting (46). Briefly, ntree was calculated by multiplying the total number of OTUs included in the analysis by a ratio of the quantity of samples in each classification category. Additionally, mtry was defined as the square root of the number of OTUs. The informative cutoff for Mean Decease Accuracy (MDA) values was determined by the absolute value of the lowest MDA measured (47). Testing for significant difference in OTU relative abundance following feature selection was performed using Wilcoxon signed-rank test with Benjamini–Hochberg correction.

## Acknowledgements

This work was funded by NIH 5T32AI007528 (J.L.L.), the Rackham Predoctoral Fellowship (J.L.L.), UM EDGE Student Fellowship (J.L.L.), and U01AI124255 (V.B.Y.). The funders had no role in study design, data collection or interpretation. The authors report no financial conflict of interest.

The authors would like acknowledging Judith Opp, April Cockburn and Harriet Carrington of the University of Michigan Microbial Systems Laboratory for constructing and sequencing the amplicon libraries. We would also like to thank Dr. Mary Riwes for providing protocols for splenocyte harvest and preparation. Finally, we would also like to thank Katherine Wozniak for helping to start and maintain the RAG1 knockout colony.

## Author Contributions

JLL and VBY conceived the study. JLL, KCV, and MLJ performed experiments and analyzed the data. All authors contributed to writing the manuscript and had access to all of the data.

---

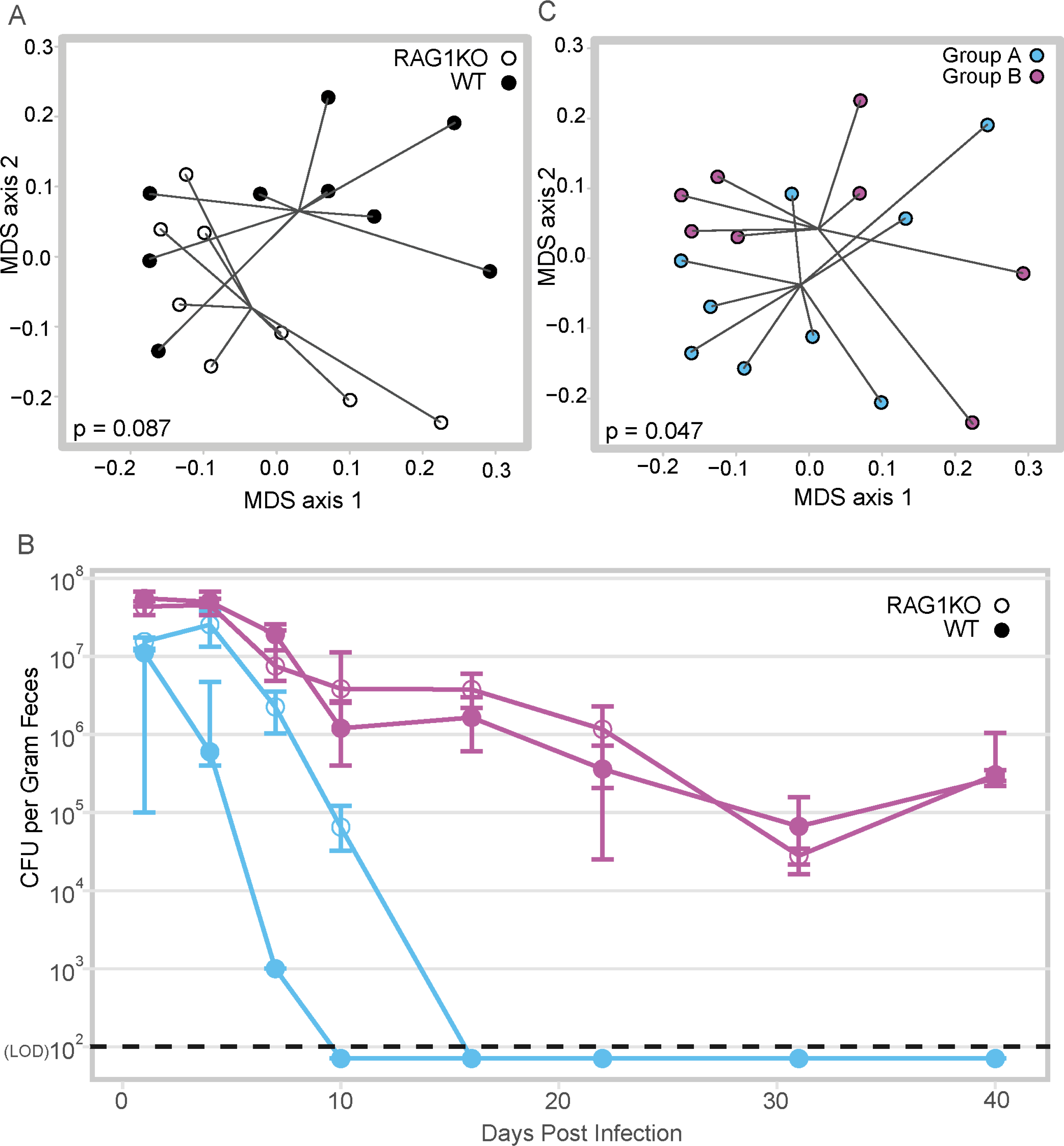

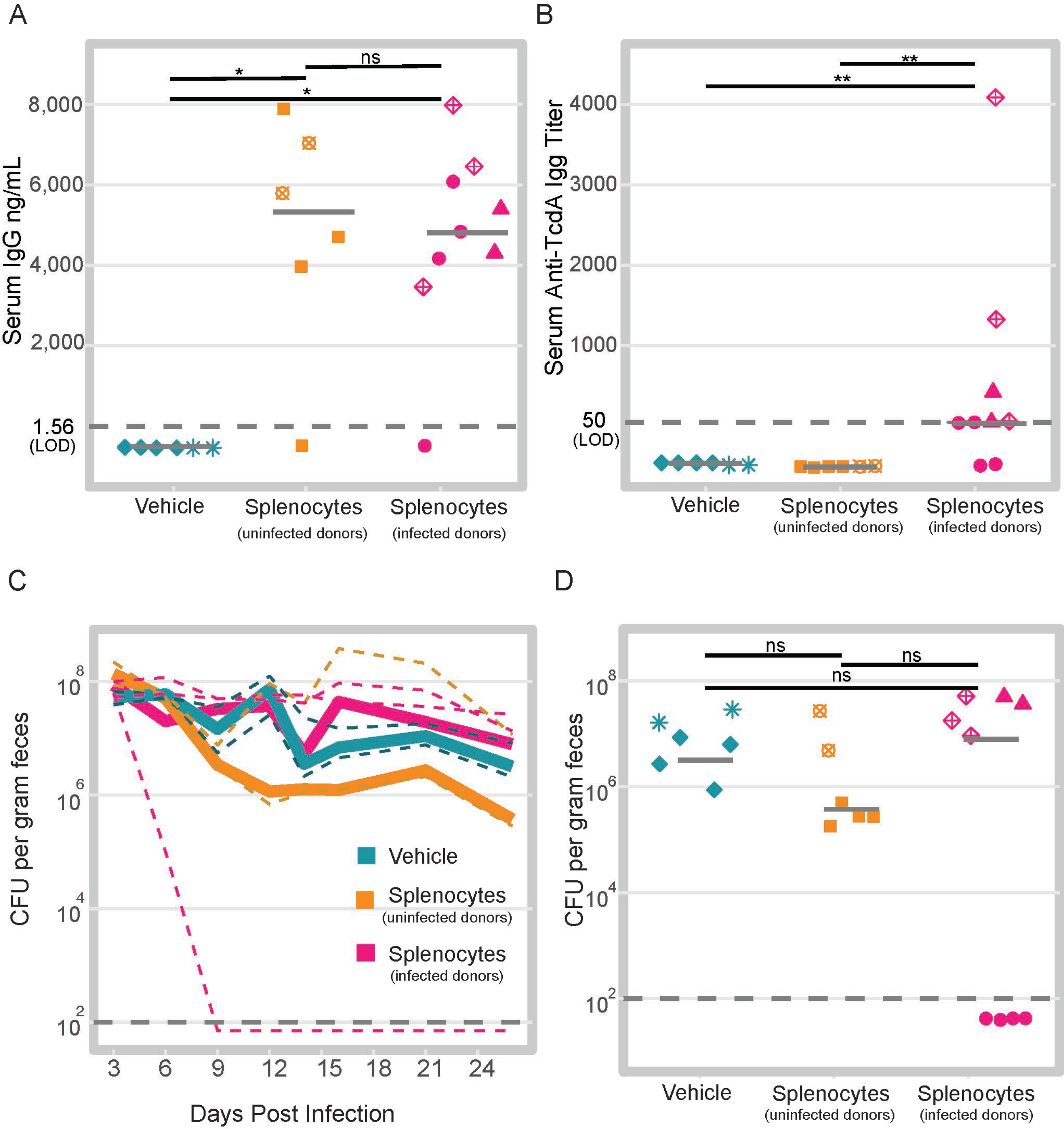

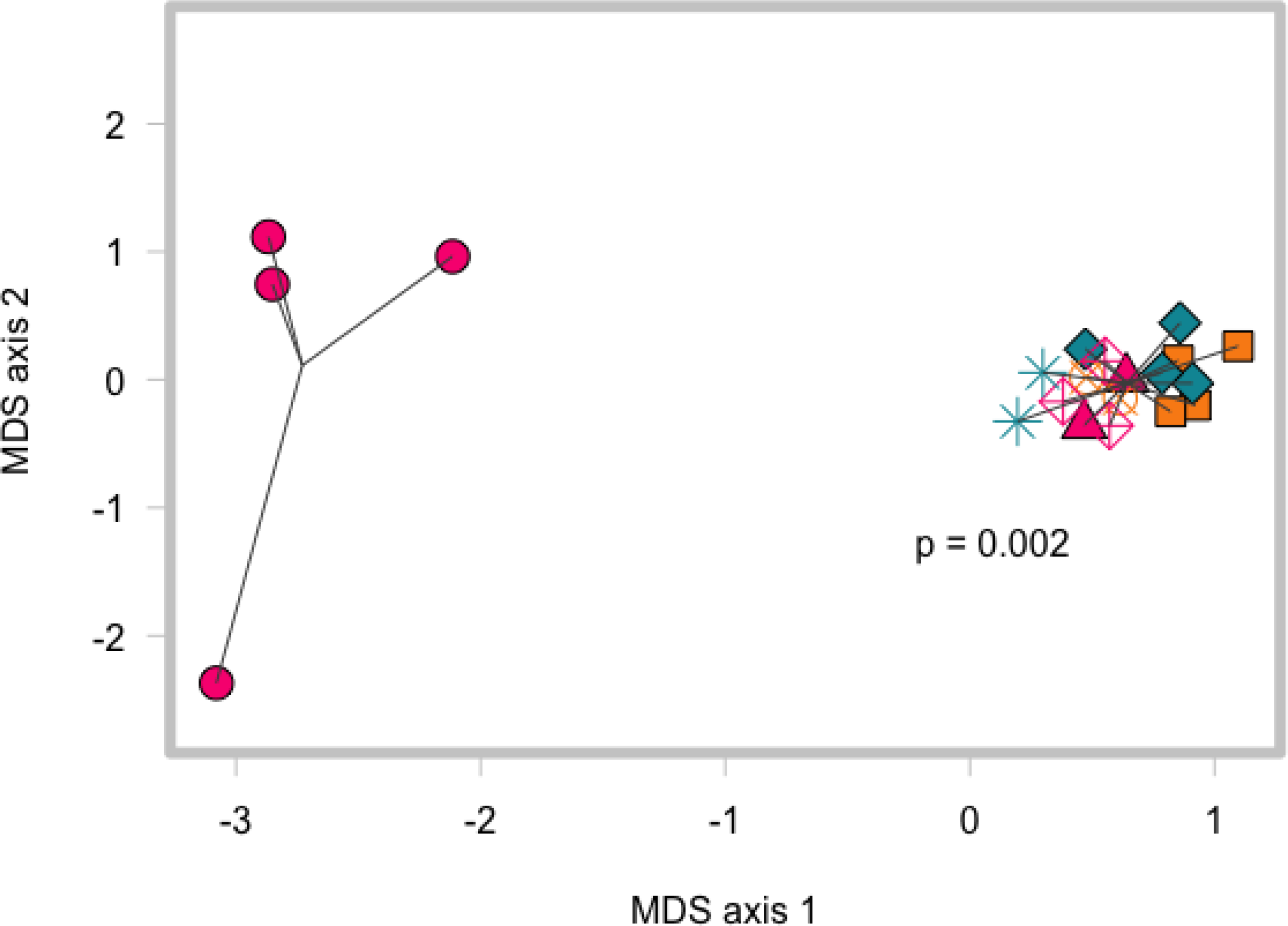

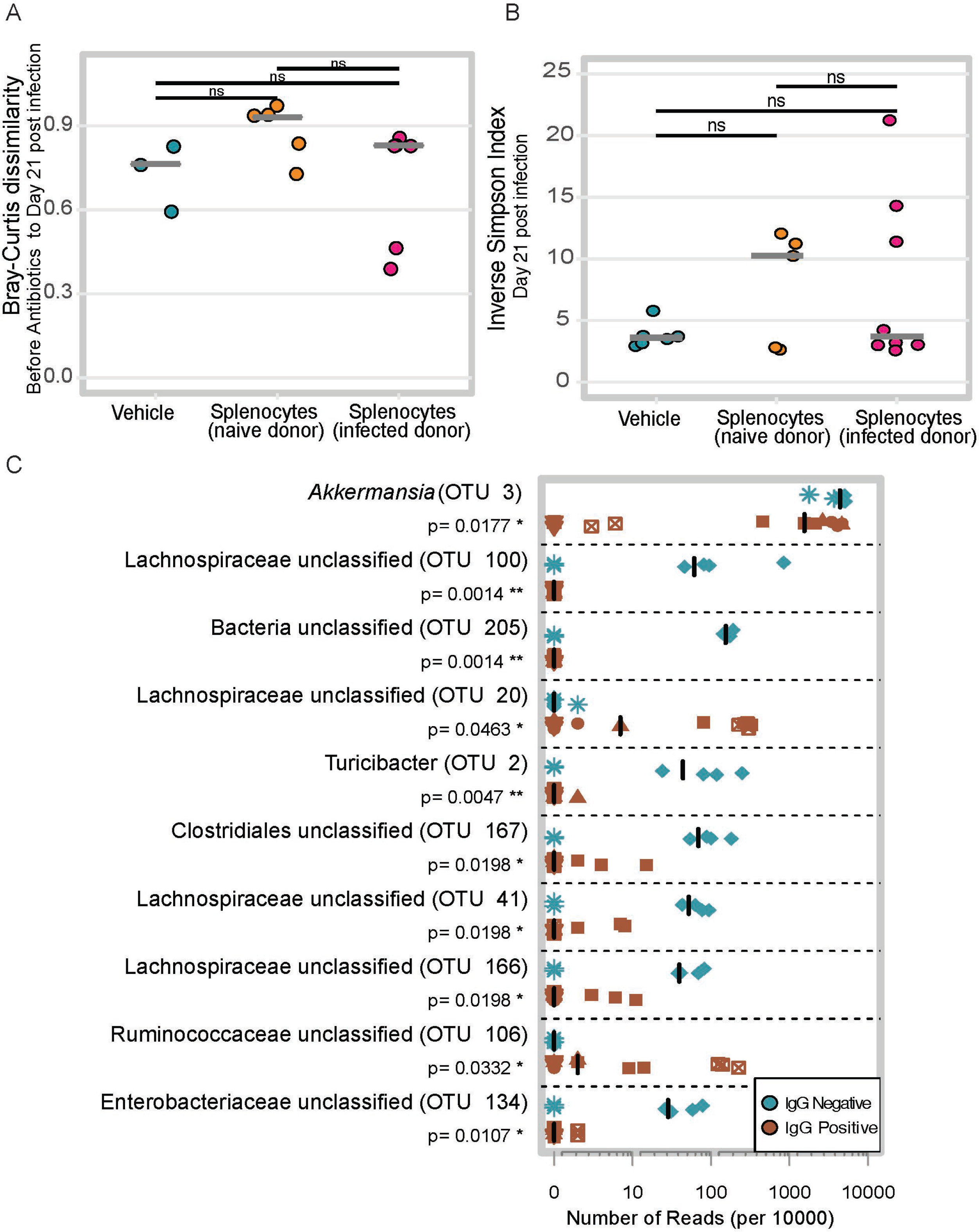

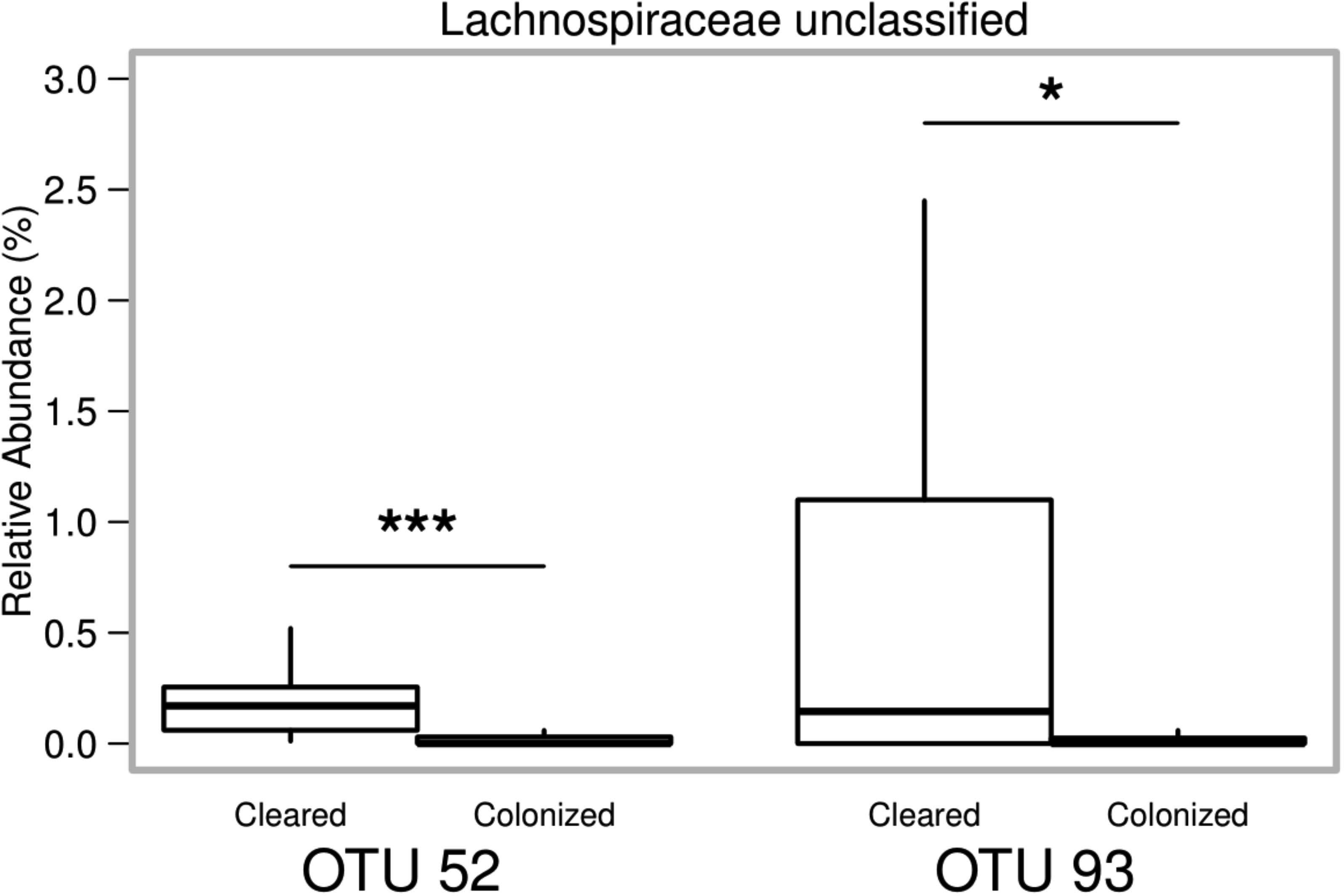

## References

1. Lessa FC, Mu Y, Bamberg WM, Beldavs ZG, Dumyati GK, Dunn JR, Farley MM, Holzbauer SM, Meek JI, Phipps EC, Wilson LE, Winston LG, Cohen JA, Limbago BM, Fridkin SK, Gerding DN, McDonald LC. 2015. Burden of Clostridium difficile infection in the United States. New England Journal of Medicine 372:825–834.

2. Chalmers JD, Akram AR, Singanayagam A, Wilcox MH, Hill AT. 2016. Risk factors for Clostridium difficile infection in hospitalized patients with community-acquired pneumonia. J Infect 73:45–53.

3. Theriot CM, Koenigsknecht MJ, Carlson PE, Hatton GE, Nelson AM, Li B, Huffnagle GB, Li J, Young VB. 2014. Antibiotic-induced shifts in the mouse gut microbiome and metabolome increase susceptibility to Clostridium difficile infection. Nature communications 5:3114–3114.

4. Carter GP, Rood JI, Lyras D. 2010. The role of toxin A and toxin B in Clostridium difficile-associated disease. Gut Microbes 1:58–64.

5. Pruitt RN, Chumbler NM, Rutherford SA, Farrow MA, Friedman DB, Spiller B, Lacy DB. 2012. Structural determinants of Clostridium difficile toxin A glucosyltransferase activity. Journal of Biological Chemistry 287:8013–8020.

6. Leslie JL, Huang S, Opp JS, Nagy MS, Kobayashi M, Young VB, Spence JR. 2015. Persistence and toxin production by Clostridium difficile within human intestinal organoids result in disruption of epithelial paracellular barrier function. Infection and Immunity 83:138–145.

7. Tam J, Beilhartz Greg L, Auger A, Gupta P, Therien Alex G, Melnyk Roman A. 2015. Small molecule inhibitors of Clostridium difficile toxin B-induced cellular damage. Chemistry … Biology 22:175–185.

8. Jiang ZD, Ajami NJ, Petrosino JF, Jun G, Hanis CL, Shah M, Hochman L, Ankoma-Sey V, DuPont AW, Wong MC, Alexander A, Ke S, DuPont HL. 2017. Randomised clinical trial: faecal microbiota transplantation for recurrent Clostridum difficile infection - fresh, or frozen, or lyophilised microbiota from a small pool of healthy donors delivered by colonoscopy. Aliment Pharmacol Ther 45:899–908.

9. Brandt LJ, Aroniadis OC, Mellow M, Kanatzar A, Kelly C, Park T, Stollman N, Rohlke F, Surawicz C. 2012. Long-Term Follow-Up of Colonoscopic Fecal Microbiota Transplant for Recurrent Clostridium difficile Infection. Am J Gastroenterol 107:1079–1087.

10. Dowle C. 2016. Faecal microbiota transplantation: a review of FMT as an alternative treatment for Clostridium difficile infection. Bioscience Horizons: The International Journal of Student Research 9:hzw007–hzw007.

11. Kyne L, Warny M, Qamar A, Kelly C. 2001. Association between antibody response to toxin A and protection against recurrent Clostridium difficile diarrhoea. Lancet 357:189–193.

12. Wilcox MH, Gerding DN, Poxton IR, Kelly C, Nathan R, Birch T, Cornely OA, Rahav G, Bouza E, Lee C, Jenkin G, Jensen W, Kim Y.S, Yoshida J, Gabryelski L, Pedley A, Eves K, Tipping R, Guris D, Kartsonis N, Dorr M.B. 2017. Bezlotoxumab for prevention of recurrent Clostridium difficile infection. New England Journal of Medicine 376:305–317.

13. Giannasca PJ, Zhang ZX, Lei WD, Boden JA, Giel MA, Monath TP, Thomas WD. 1999. Serum antitoxin antibodies mediate systemic and mucosal protection from Clostridium difficile disease in hamsters. Infection and immunity 67:527–538.

14. Johnston PF, Gerding DN, Knight KL. 2014. Protection from Clostridium difficile infection in CD4 T cell− and polymeric immunoglobulin receptor-deficient mice. Infection and Immunity 82:522–531.

15. Bruxelle JF, Mizrahi A, Hoys S, Collignon A, Janoir C, Péchiné S. 2016. Immunogenic properties of the surface layer precursor of Clostridium difficile and vaccination assays in animal models. Anaerobe 37:78–84.

16. Ghose C, Eugenis I, Sun X, Edwards AN, McBride SM, Pride DT, Kelly CP, Ho DD. 2016. Immunogenicity and protective efficacy of recombinant Clostridium difficile flagellar protein FliC. Emerging Microbes … Infections 5:e8.

17. Lawley TD, Clare S, Walker AW, Stares MD, Connor TR, Raisen C, Goulding D, Rad R, Schreiber F, Brandt C, Deakin LJ, Pickard DJ, Duncan SH, Flint HJ, Clark TG, Parkhill J, Dougan G. 2012. Targeted restoration of the intestinal microbiota with a simple, defined bacteriotherapy resolves relapsing Clostridium difficile disease in mice. PLoS Pathog 8:e1002995.

18. Round JL, Mazmanian SK. 2009. The gut microbiome shapes intestinal immune responses during health and disease. Nature reviews Immunology 9:313–323.

19. Rooks MG, Garrett WS. 2016. Gut microbiota, metabolites and host immunity. Nat Rev Immunol 16:341–352.

20. Theriot CM, Bowman AA, Young VB. 2016. Antibiotic-induced alterations of the gut microbiota alter secondary bile acid production and allow for Clostridium difficile spore germination and outgrowth in the large intestine. mSphere 1.

21. Zhang H, Sparks JB, Karyala SV, Settlage R, Luo XM. 2015. Host adaptive immunity alters gut microbiota. ISME J 9:770–781.

22. Kamada N, Sakamoto K, Seo SU, Zeng MY, Kim YG, Cascalho M, Vallance BA, Puente JL, Nunez G. 2015. Humoral Immunity in the Gut Selectively Targets Phenotypically Virulent Attaching-and-Effacing Bacteria for Intraluminal Elimination. Cell Host Microbe 17:617–27.

23. Kamada N, Kim Y.G, Sham H, Vallance B, Puente J, Martens E, Núñez G. 2012. Regulated virulence controls the ability of a pathogen to compete with the gut microbiota. Science (New York, NY) 336:1325–1329.

24. McDermott AJ, Huffnagle GB. 2014. The microbiome and regulation of mucosal immunity. Immunology 142:24–31.

25. Plovier H, Everard A, Druart C, Depommier C, Van Hul M, Geurts L, Chilloux J, Ottman N, Duparc T, Lichtenstein L, Myridakis A, Delzenne NM, Klievink J, Bhattacharjee A, van der.Ark KCH, Aalvink S, Martinez LO, Dumas M.E, Maiter D, Loumaye A, Hermans MP, Thissen J.P, Belzer C, de Vos WM, Cani PD. 2017. A purified membrane protein from Akkermansia muciniphila or the pasteurized bacterium improves metabolism in obese and diabetic mice. Nat Med 23:107–113.

26. Everard A, Belzer C, Geurts L, Ouwerkerk JP, Druart C, Bindels LB, Guiot Y, Derrien M, Muccioli GG, Delzenne NM, de Vos WM, Cani PD. 2013. Cross-talk between Akkermansia muciniphila and intestinal epithelium controls diet-induced obesity. Proc Natl Acad Sci U S A 110:9066–71.

27. Schubert AM, Sinani H, Schloss PD. 2015. Antibiotic-induced alterations of the murine gut microbiota and subsequent effects on colonization resistance against Clostridium difficile. mBio 6.

28. Reeves AE, Theriot CM, Bergin IL, Huffnagle GB, Schloss PD, Young VB. 2010. The interplay between microbiome dynamics and pathogen dynamics in a murine model of Clostridium difficile Infection. Gut microbes 2:145–158.

29. Reeves AE, Koenigsknecht MJ, Bergin IL, Young VB. 2012. Suppression of Clostridium difficile in the gastrointestinal tracts of germfree mice inoculated with a murine isolate from the family Lachnospiraceae. Infection and immunity 80:3786–3794.

30. Stappenbeck TS, Virgin HW. 2016. Accounting for reciprocal host–microbiome interactions in experimental science. Nature 534:191–199.

31. Perez J, Springthorpe, V. S. … Sattar, S. A.. 2011. Clospore: a liquid medium for producing high titers of semi-purified spores of Clostridium difficile. J AOAC Int 94:618–626.

32. George WL, Sutter VL, Citron D, Finegold SM. 1979. Selective and differential medium for isolation of Clostridium difficile. Journal of Clinical Microbiology 9:214–219.

33. Trindade BC, Theriot CM, Leslie JL, Carlson Jr PE, Bergin IL, Peters-Golden M, Young VB, Aronoff DM. 2014. Clostridium difficile-induced colitis in mice is independent of leukotrienes. Anaerobe 30:90–98.

34. Seekatz AM, Theriot CM, Molloy CT, Wozniak KL, Bergin IL, Young VB. 2015. Fecal Microbiota Transplantation Eliminates Clostridium difficile in a Murine Model of Relapsing Disease. Infect Immun 83:3838–46.

35. Kozich JJ, Westcott SL, Baxter NT, Highlander SK, Schloss PD. 2013. Development of a dual-index sequencing strategy and curation pipeline for analyzing amplicon sequence data on the MiSeq Illumina sequencing platform. Applied and Environmental Microbiology 79:5112–5120.

36. Schloss PD, Westcott SL, Ryabin T, Hall JR, Hartmann M, Hollister EB, Lesniewski RA, Oakley BB, Parks DH, Robinson CJ, Sahl JW, Stres B, Thallinger GG, Van Horn DJ, Weber CF. 2009. Introducing mothur: Open-Source, Platform-Independent, Community-Supported Software for Describing and Comparing Microbial Communities. Applied and Environmental Microbiology 75:7537–7541.

37. Quast C, Pruesse E, Yilmaz P, Gerken J, Schweer T, Yarza P, Peplies J, Glöckner FO. 2013. The SILVA ribosomal RNA gene database project: improved data processing and web-based tools. Nucleic Acids Research 41:D590–D596.

38. Edgar RC, Haas BJ, Clemente JC, Quince C, Knight R. 2011. UCHIME improves sensitivity and speed of chimera detection. Bioinformatics 27:2194–2200.

39. Wang Q, Garrity GM, Tiedje JM, Cole JR. 2007. Naive Bayesian classifier for rapid assignment of rRNA sequences into the new bacterial taxonomy. Appl Environ Microbiol 73:5261–7.

40. Westcott SL, Schloss PD. 2017. OptiClust, an improved method for assigning amplicon-based sequence data to operational taxonomic units. mSphere 2.

41. Segata N, Izard J, Waldron L, Gevers D, Miropolsky L, Garrett WS, Huttenhower C. 2011. Metagenomic biomarker discovery and explanation. Genome Biology 12:R60.

42. R Core Team. 2017. R: A Language and Environment for Statistical Computing, R Foundation for Statistical Computing, Vienna, Austria. http://www.R-project.org/

43. Jari Oksanen FGB, Michael Friendly, Roeland Kindt, Pierre Legendre, Dan McGlinn, Peter R. Minchin, R. B. O’Hara, Gavin L. Simpson, Peter Solymos, M. Henry H. Stevens, Eduard Szoecs.and Helene Wagner. 2017. vegan: Community Ecology Package., vR package version 2.4–3. CRAN. https://CRAN.R-project.org/package=vegan.

44. Breiman L. 2001. Random Forests. Machine Learning 45:5–32.

45. Liaw A, Wiener M. 2002. Classification and regression by randomForest. R news 2:18–22.

46. Huang BFF, Boutros PC. 2016. The parameter sensitivity of random forests. BMC Bioinformatics 17:331.

47. Strobl C, Malley J, Tutz G. 2009. An introduction to recursive partitioning: rationale, application and characteristics of classification and regression trees, bagging and random forests. Psychological methods 14:323–348.

